# Comprehensive genome data analysis establishes a triple whammy of carbapenemases, ICEs and multiple clinically-relevant bacteria

**DOI:** 10.1101/678748

**Authors:** João Botelho, Joana Mourão, Adam P. Roberts, Luísa Peixe

## Abstract

Carbapenemases inactivate most β-lactam antibiotics, including carbapenems and have been frequently reported among *Enterobacteriaceae*, *Acinetobacter* spp. and *Pseudomonas* spp. Traditionally, the horizontal gene transfer of carbapenemase encoding genes (CEGs) has been linked to plasmids. However, given that integrative and conjugative elements (ICEs) are possibly the most abundant conjugative elements among prokaryotes, we conducted an *in-silico* analysis to ascertain the likely role of ICEs in the spread of CEGs among all bacterial genomes (n=182,663). We detected 17,520 CEGs, of which 66 were located within putative ICEs among several bacterial species (including clinically-relevant bacteria as *Pseudomonas aeruginosa*, *Klebsiella pneumoniae* and *Escherichia coli*). Most CEGs detected within ICEs belong to the IMP, NDM and SPM metallo-beta-lactamase families, and the serine beta-lactamase KPC and GES families. Different mechanisms were likely responsible for acquisition of these genes. The majority of CEG-bearing ICEs belong to the MPF_G_, MPF_T_ and MPF_F_ classes and often encode resistance to other antibiotics (e.g., aminoglycosides and fluoroquinolones). This study provides a snapshot of the different CEGs associated with ICEs among available bacterial genomes and sheds light on the underappreciated contribution of ICEs to the spread of carbapenem resistance globally.

**Author Notes:** All supporting data has been provided within the article or through supplementary data files. Supplementary material is available with the online version of this article.

**Impact Statement:** Carbapenems are commonly used to treat severe infections in humans. Resistance is often mediated by carbapenemases. These enzymes degrade carbapenems and are frequently present in plasmids. Here, we demonstrate that common carbapenemase-encoding genes (CEGs) found in clinical isolates (e.g. *bla*_KPC_, *bla*_GES_, *bla*_IMP_, *bla*_NDM_, *bla*_VIM_) can also be located within integrative and conjugative elements (ICEs). CEG-bearing ICEs belong to three mating-pair formation families. These mobile elements may be particularly important in bacteria where plasmids do not seem to play a significant role in the spread of antibiotic resistance genes, as *Pseudomonas* spp. This study considerably expands the knowledge of the repertoire of CEGs-bearing ICEs among clinically-relevant bacterial pathogens, such as *Pseudomonas aeruginosa*, *Klebsiella pneumoniae* and *Escherichia coli*.

**Data Summary:** All the bacterial genomes scanned in this study have been deposited previously in the National Center for Biotechnology Information genome database and are listed on the supplementary tables. The extracted 66 ICEs in fasta format and the outputs for the profile HMMs scanned on the 386 putative MGEs identified in this study are deposited on figshare at https://figshare.com/projects/_Comprehensive_genome_data_analysis_establishes_a_triple_whammy_of_carbapenemases_ICEs_and_multiple_clinically-relevant_bacteria/78369.

## Introduction

Due to the importance of carbapenems for the treatment of severe infections in humans, the WHO stated that these antibiotics should be reserved for infections caused by multidrug resistant Gram-negative bacteria in humans [1]. Recently, the same agency presented a list of bacterial pathogens for which research and development of new antibiotics are urgently required, and the top priority pathogens were the carbapenem-resistant strains of *Acinetobacter baumannii*, *Pseudomonas aeruginosa* and *Enterobacteriaceae* [2].

The evolution of carbapenem-resistance in bacteria is often driven by the horizontal gene transfer (HGT) of carbapenemase-encoding genes (CEGs) [3, 4]. Carbapenemases are beta-lactamases able to hydrolyse carbapenems as well as most of other beta-lactam antibiotics. These enzymes are members of the serine beta-lactamases classes A and D, and the class B metallo-beta-lactamases [5]. The CEGs are often located on integrons or transposons that themselves target mobile genetic elements (MGEs) such as plasmids [3, 4], which makes the dissemination of these genes unpredictably complex within bacterial communities. Recently, it was demonstrated that another type of MGE, the integrative and conjugative elements (ICEs), are likely to play a significant role as vehicles for the dissemination of CEGs among *P. aeruginosa* [6]. Besides genes conferring antibiotic resistance, ICEs may harbour additional cargo genes that provide an adaptive advantage over other elements. Some of these examples include the presence of the siderophore yersiniabactin encoded within the ICEKp in hypervirulent clonal-group CG23 *Klebsiella pneumoniae* [7]; the Tn*5252*-related ICEs carrying bacteriocin clusters in *Streptococcus suis* [8]; the type I-C CRISPR-Cas systems identified within pKLC102-like ICEs in *P. aeruginosa* [9] and the type III restriction-modification systems from SXT/R391-related ICEs in *Shewanella* spp. [10].

ICEs are self-transmissible MGEs that can integrate into and excise from the genome (as transposons and phages do) and can exist as circular, sometimes replicable, extrachromosomal elements and be transferred by conjugation (as some plasmids do) [11–14]. ICE*clc* from *Pseudomonas knackmussii* [15], SXT from *Vibrio cholerae* [16], pKLC102 from *P. aeruginosa* [17], and Tn*4371* from *Ralstonia oxalatica* [18] are among the most well studied ICEs. ICEs appear to have a bipartite lifestyle that shifts between vertical and horizontal transmission [12, 19, 20]. HGT by conjugation requires three main components: a relaxase (MOB), a mating pair formation (MPF) system, and a type-IV coupling protein, with the last two forming a spanning-membrane multi-protein complex named type-IV secretion system (T4SS) [21]. Until now, eight MPF classes were proposed (B, C, F, FA, FATA, G, I and T), based on the phylogeny of VirB4, the only ubiquitous protein among the T4SS. The MPF_T_ is widely distributed both in conjugative plasmids and ICEs, while MPF_F_ is more prevalent in plasmids and MPF_G_ on ICEs [11].

Given that ICEs were identified in most bacterial clades and were proposed to be more prevalent than conjugative plasmids [11], we conducted an *in silico* analysis to explore the distribution of CEG-bearing ICEs among all sequenced bacterial genomes. Our results demonstrate that CEG-bearing ICEs belong to three MPF families and are primarily located in several clinically-relevant bacterial pathogens. Our analysis highlights the importance of thoroughly investigating these elements as important vehicles for the spread of antibiotic resistance (AR), particularly to carbapenems.

## Material and methods

### Bacterial genomes and carbapenemases search

In **Figure 1**, we present the workflow used in this study, from the acquisition of bacterial genomes to the identification and characterization of putative ICEs. We retrieved all bacterial genomes available in NCBI Reference Sequence Database (RefSeq, accessed on 21/03/2020), including complete and draft genome sequences, using ncbi-genome-download v0.2.12 (https://github.com/kblin/ncbi-genome-download). We downloaded over 6,000 curated AR protein sequences from AMRfinder database (https://ftp.ncbi.nlm.nih.gov/pathogen/Antimicrobial_resistance/AMRFinderPlus/database/3.6/2020-01-22.1/) [22] and we built an in-house database only including the proteins that code for a carbapenemase (n=1014, **Table S1**). We then blasted the genomes against the extracted carbapenemases using diamond v0.9.29.130 (http://www.diamondsearch.org/index.php) [23], using minimum 100% identity and subject cover and with the sensitive mode enabled.

**Figure 1.**
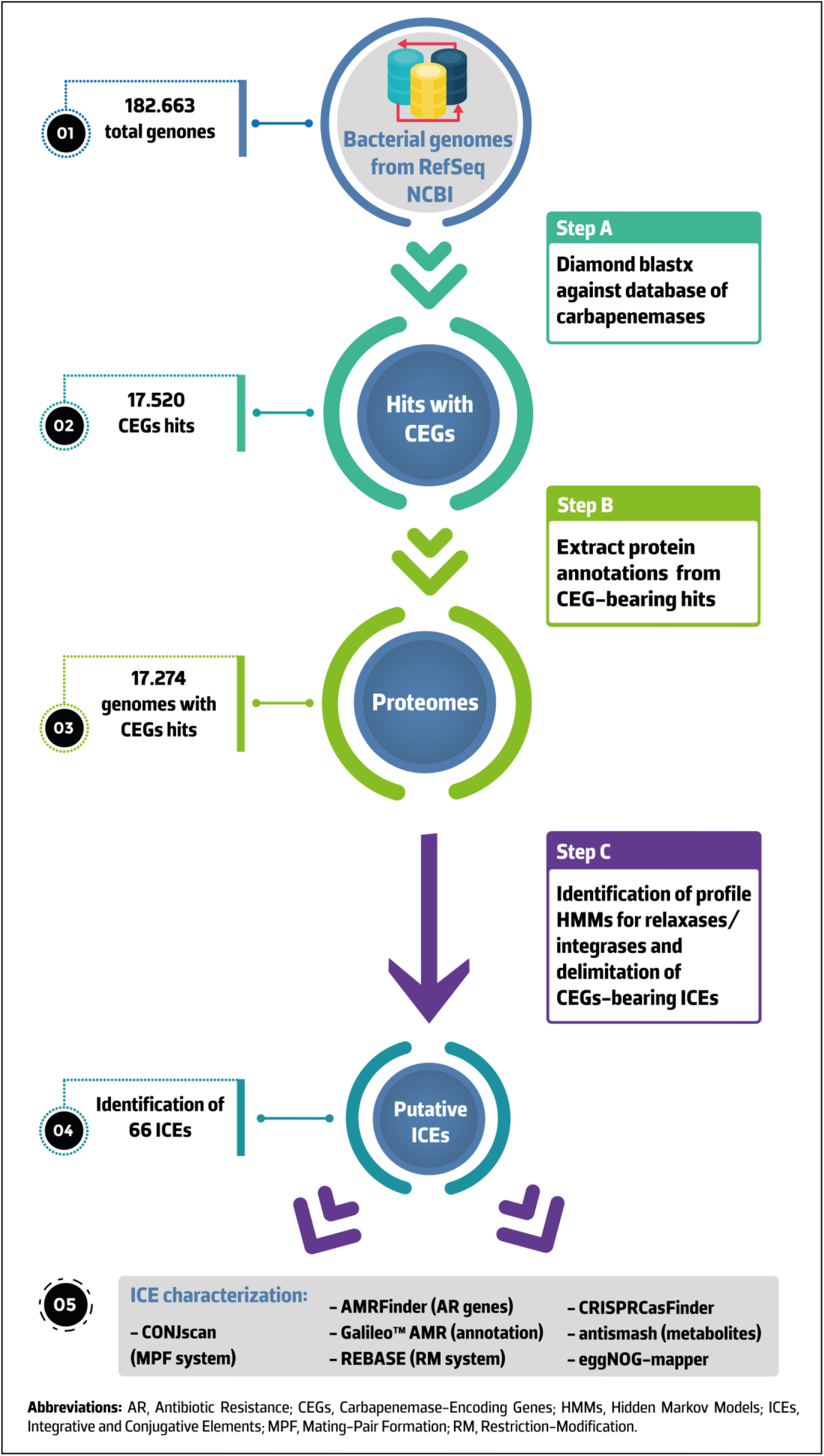
Overview of the workflow followed in this study. All assemblies available in NCBI RefSeq were downloaded and blasted against an in-house database of carbapenemases using diamond blastx (Step 1). NCBI annotated proteins from CGE-bearing genomes were then extracted (Step 2) and used for the identification of relaxase and serine or tyrosine recombinase (Step 3). Search of directed repeats and delimitation of putative ICEs was also performed. CONJscan was used to identify the MPF family of each element. We then looked for AR genes, restriction-modification systems, CRISPR arrays and their associated (Cas) proteins as well as secondary metabolites within extracted ICEs. We also characterized the functional annotations of their proteomes and the MLST of the genomes carrying a CEG-bearing ICE.

### Tracing ICEs among the bacterial genomes

The RefSeq protein files from the CEG-bearing genomes identified by diamond were extracted. We used the hmmsearch function of the HMMER3 software package v3.3 (http://hmmer.org/) [24] to search the proteomes against the standalone version of MOBfamDB, a curated hidden Markov models (HMM) relaxase database (https://castillo.dicom.unican.es/mobscan_about/) [25]. We also used this function to search the pfam v33.0 database for tyrosine or serine recombinase accessions numbers (Pfam IDs PF00589 and PF07508). The hmmsearch command was used with default parameters and an E-value threshold of 0.01. The CEG-bearing genomes with relaxase and integrase hits were further analysed. We used the Find Repeats tool from Geneious Prime^®^ 2020.0.4 (https://www.geneious.com) to inspect the hits for direct repeats. To delimit CEG-harbouring ICEs, we manually scanned candidate terminal regions with direct repetitions of the 3’ end from tRNA genes located next to the integrase-encoding gene. When no tRNA gene was identified next to this gene, we scanned the presence of direct repeats next to the integrase-encoding gene and next to candidate terminal regions. To assist in identifying putative terminal regions, we looked for blocks of DNA with variation in GC content. To predict the MPF families, the translated coding sequences of delimited ICEs were analysed on the standalone CONJscan module of MacSyFinder v1.0.5 (https://github.com/gem-pasteur/macsyfinder) [26, 27]. To identify the multi-locus sequence type of the genomes containing CEG-bearing ICEs, we used mlst v2.16.1 (https://github.com/tseemann/mlst), which scans the genomes against PubMLST typing schemes (https://pubmlst.org/) [28].

### Characterization of the CEG-bearing ICEs

Screening of AR genes among ICEs was performed using amrfinder v3.6.10 (https://github.com/ncbi/amr) [22]. The genetic platforms involved in the acquisition of CEGs by ICEs were annotated using Galileo™ AMR (https://galileoamr.arcbio.com/mara/) (Arc Bio, Cambridge, MA) [29]. We ran our extracted ICEs against REBASE (http://rebase.neb.com/rebase/rebase.html) to look for restriction-modification systems [30]. We used CRISPRCasFinder (https://crisprcas.i2bc.paris-saclay.fr/CrisprCasFinder/Index) to look for CRISPR (clustered regularly interspaced short palindromic repeats) arrays and their associated (Cas) proteins within ICE sequences [31]. Secondary metabolite biosynthetic gene clusters were traced using antismash v5.1.2 (https://antismash.secondarymetabolites.org/#!/start) [32]. We used eggNOG-mapper v2 (http://eggnog-mapper.embl.de/) for functional annotation based on orthology assignments of the ICE proteomes [33].

## Results

### Carbapenemase-encoding genes are mainly found in Proteobacteria

We retrieved a total of 182,663 bacterial genomes from NCBI (16,798 complete genomes and 165,865 genomes assembled at the chromosome, scaffold or contig level). We identified a total of 17,520 CEGs, with 1,422 CEGs on 1,236 complete genomes (including 512 chromosomes and 724 plasmids) and 16,098 CEGs on 16,038 draft genomes (**Table S2**). We identified a total of 377 carbapenemase variants among the 17,520 hits. Our results show that CEGs are mostly located on Proteobacteria and dominated by clinically-relevant pathogens as *A. baumannii*, *K. pneumoniae*, *Escherichia coli* and *P. aeruginosa* (**Table S2**). These genomes encode a wide diversity of carbapenemases, including OXA-23 (15.6%, n=2,739/17,520), KPC-2 (13.2%), KPC-3 (8.9%) and NDM-1 (6.7%). As we are tracing MGEs integrated in the chromosome, the 724 plasmid hits in complete genomes were excluded from the analysis. Among the hits on the 16,038 draft genomes, we filtered out sequences with the word ‘plasmid’ present on the fasta-header (n=131, **Table S3**). To maximise the chances of detecting entire ICEs, we filtered out sequences shorter than 40kb (n=10,050). All the excluded ones are available for analysis in **Table S4**. The remaining sequences from draft genomes (n=5,857) were inspected for the presence of CEG-harbouring ICEs.

### A large proportion of CEG-bearing ICEs belong to three families and target clinically-relevant gram-negative bacteria

We identified a total of 66 putative ICEs, including 42 newly characterized elements associated with 17 different CEGs (**Table 1** and **Figure 2**). We could predict the boundaries from 55 of these elements (**Table S5**). The terminal region of the remaining 11 putative ICEs could not be determined due to a fragmented contig or assembly gaps within the sequence. Nearly half of the putative ICEs (48.5%, n=32/66) were integrated at the 3’ end of a tRNA^Gly^ gene. Integration next to random genes was also observed (**Figure 3** and **Table S5**). The bacterial hosts housing these elements belong to 23 sequence types (STs) (**Table S5**).

**Table 1.**
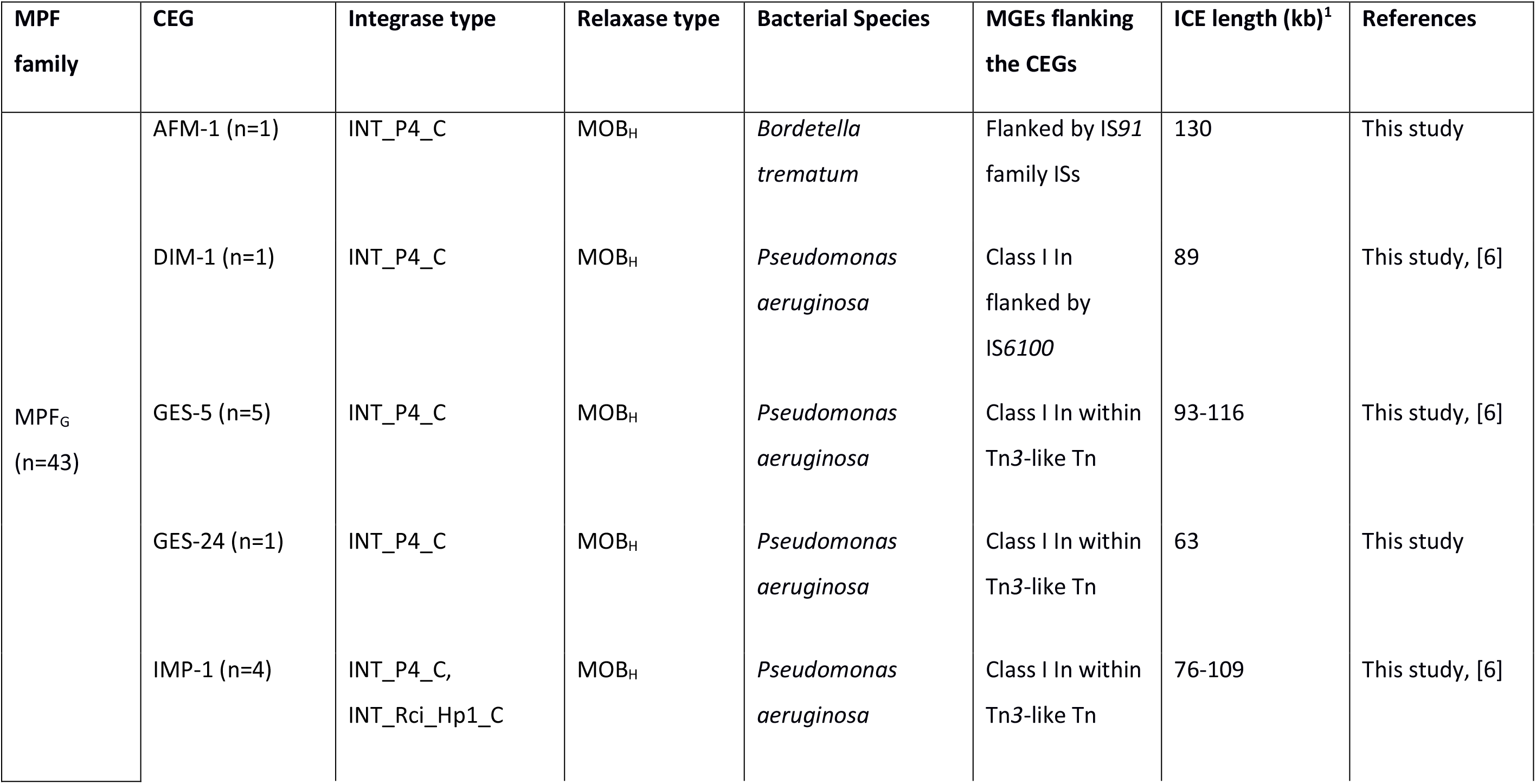
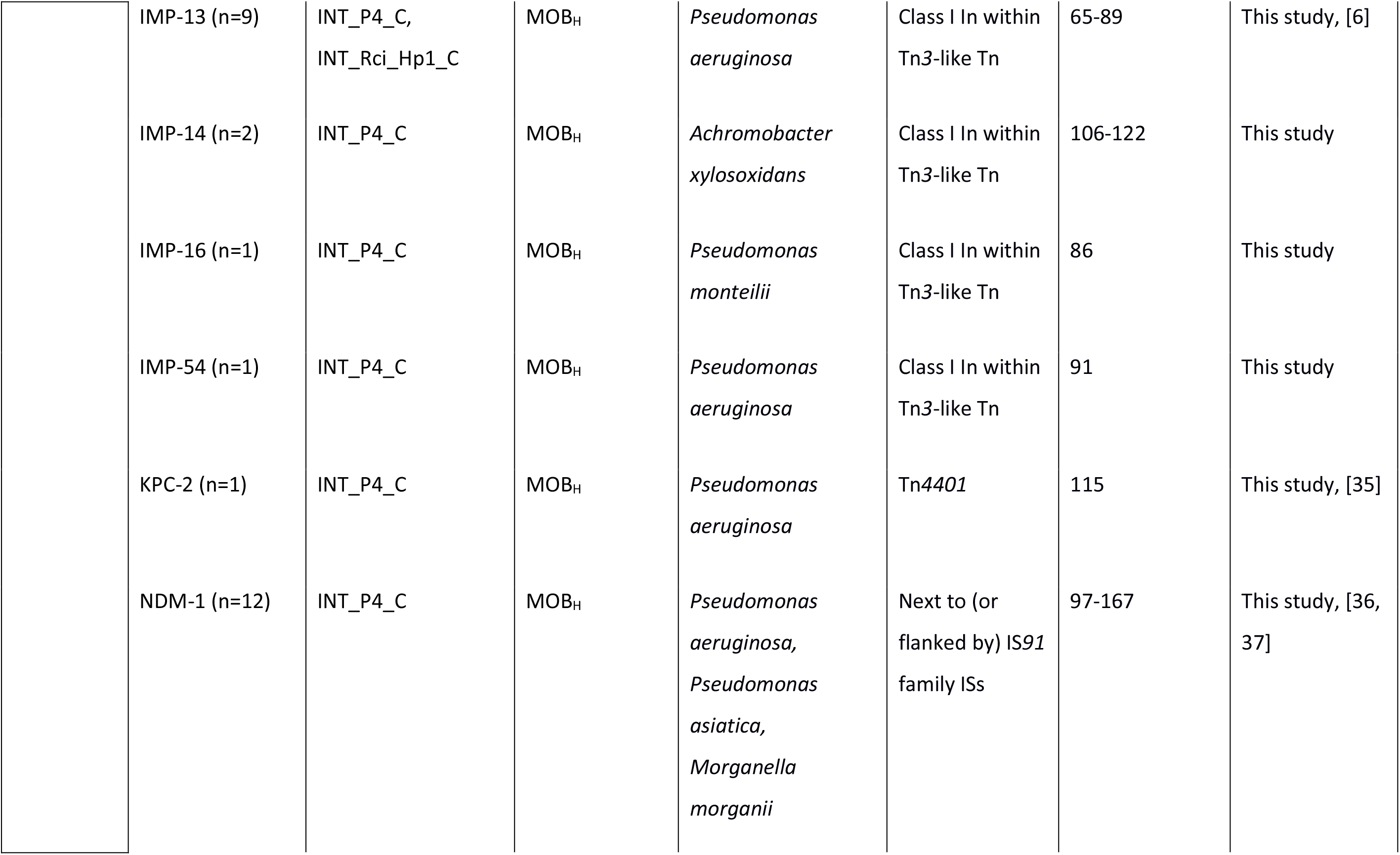
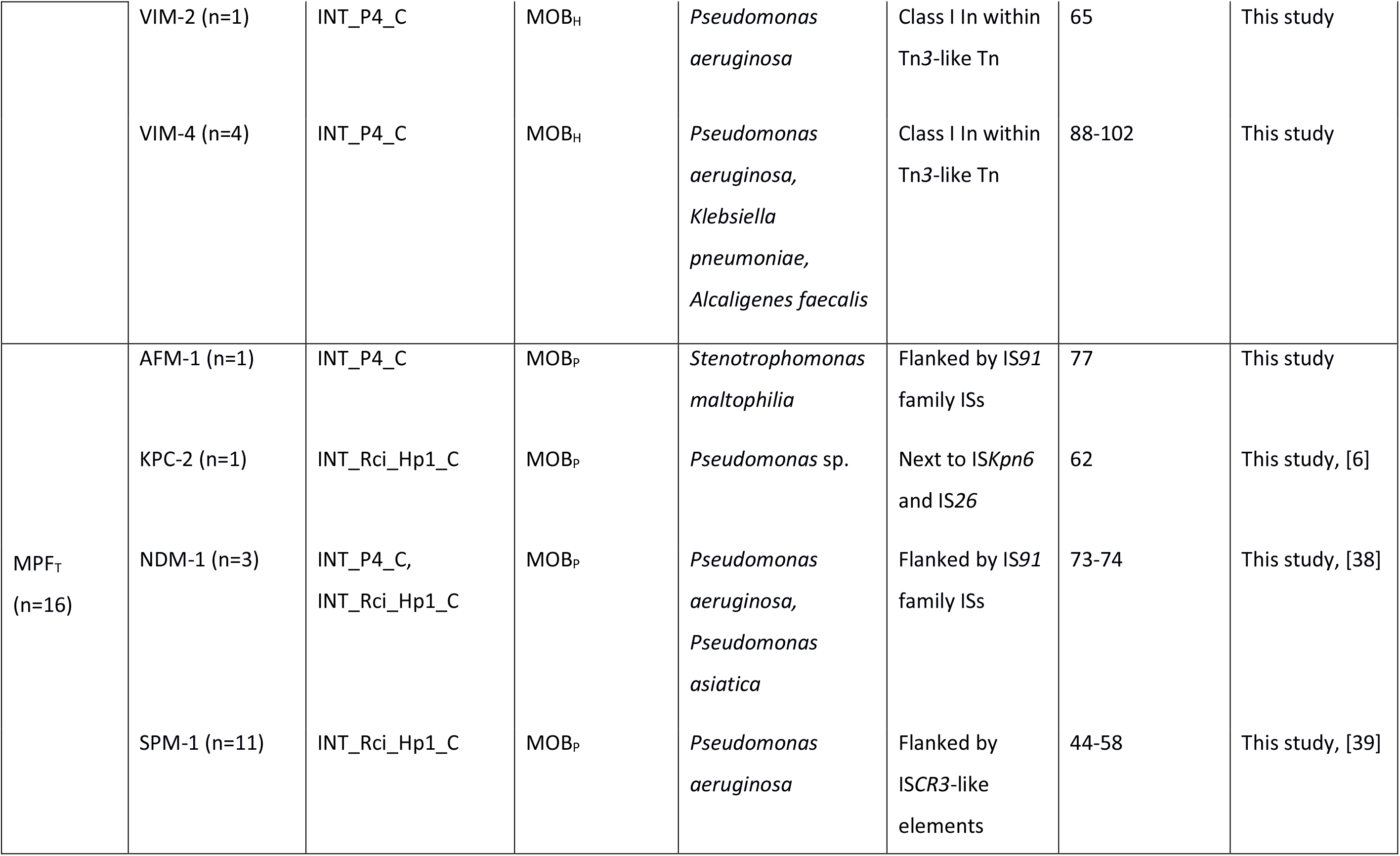
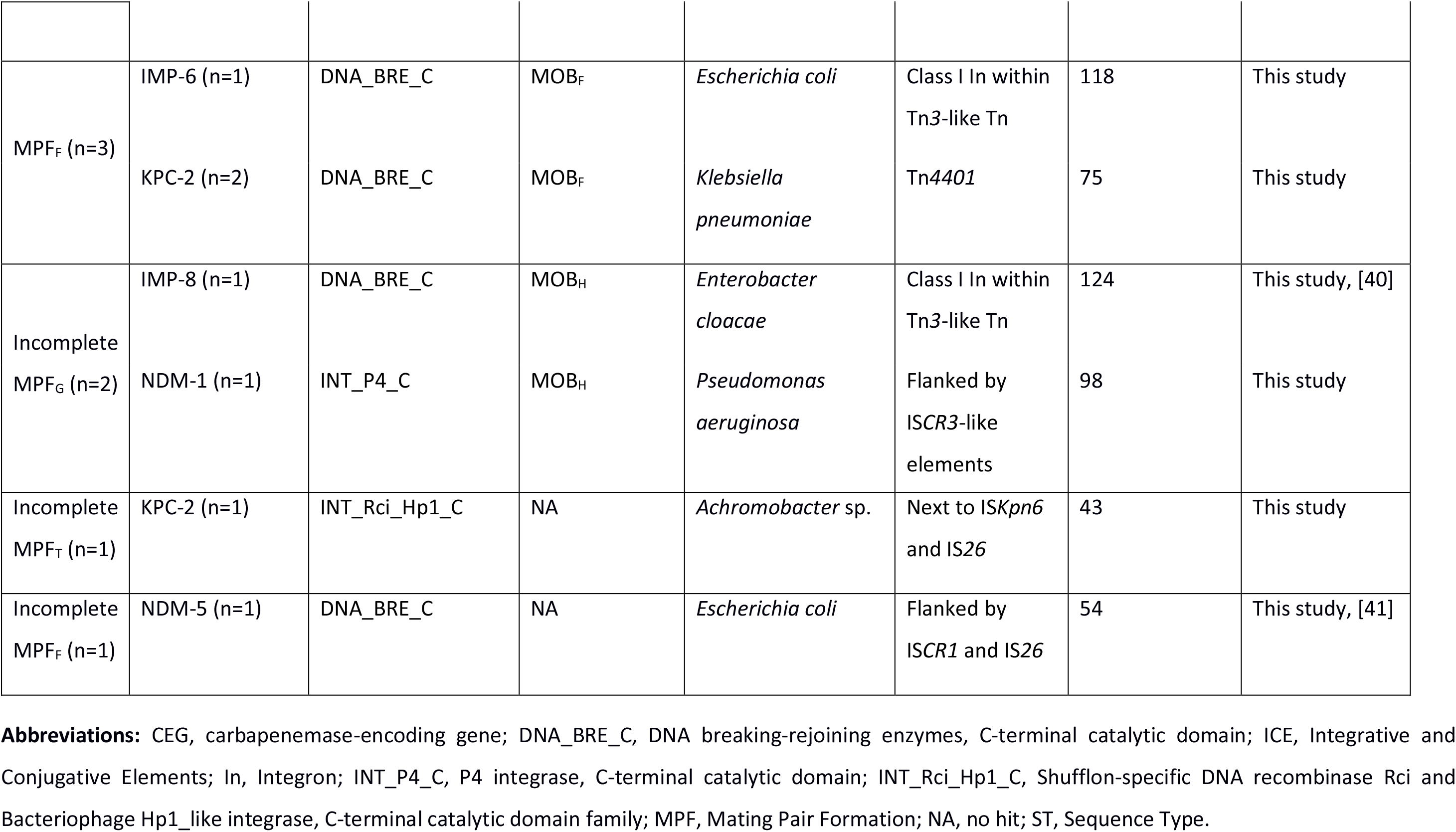
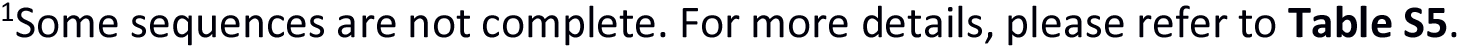
Diversity and characterization of carbapenemases-encoding genes in integrative and conjugative elements-associated genomes.

| MPF family | CEG | Integrase type | Relaxase type | Bacterial Species | MGEs flanking the CEGs | ICE length (kb)<sup>1</sup> | References |
| --- | --- | --- | --- | --- | --- | --- | --- |
| MPF<sub>G</sub><br>(n=43) | AFM-1 (n=1) | INT_P4_C | MOB<sub>H</sub> | *Bordetella trematum* | Flanked by IS91 family ISs | 130 | This study |
|  | DIM-1 (n=1) | INT_P4_C | MOB<sub>H</sub> | *Pseudomonas aeruginosa* | Class I In flanked by IS6100 | 89 | This study, [6] |
|  | GES-5 (n=5) | INT_P4_C | MOB<sub>H</sub> | *Pseudomonas aeruginosa* | Class I In within Tn3-like Tn | 93-116 | This study, [6] |
|  | GES-24 (n=1) | INT_P4_C | MOB<sub>H</sub> | *Pseudomonas aeruginosa* | Class I In within Tn3-like Tn | 63 | This study |
|  | IMP-1 (n=4) | INT_P4_C,<br>INT_Rci_Hp1_C | MOB<sub>H</sub> | *Pseudomonas aeruginosa* | Class I In within Tn3-like Tn | 76-109 | This study, [6] |
|  | IMP-13 (n=9) | INT_P4_C,<br>INT_Rci_Hp1_C | MOB<sub>H</sub> | *Pseudomonas aeruginosa* | Class I In within<br>Tn3-like Tn | 65-89 | This study, [6] |
|  | IMP-14 (n=2) | INT_P4_C | MOB<sub>H</sub> | *Achromobacter xylosoxidans* | Class I In within<br>Tn3-like Tn | 106-122 | This study |
|  | IMP-16 (n=1) | INT_P4_C | MOB<sub>H</sub> | *Pseudomonas monteilii* | Class I In within<br>Tn3-like Tn | 86 | This study |
|  | IMP-54 (n=1) | INT_P4_C | MOB<sub>H</sub> | *Pseudomonas aeruginosa* | Class I In within<br>Tn3-like Tn | 91 | This study |
|  | KPC-2 (n=1) | INT_P4_C | MOB<sub>H</sub> | *Pseudomonas aeruginosa* | Tn4401 | 115 | This study, [35] |
|  | NDM-1 (n=12) | INT_P4_C | MOB<sub>H</sub> | *Pseudomonas aeruginosa*,<br>*Pseudomonas asiatica*,<br>*Morganella morganii* | Next to (or<br>flanked by) IS91<br>family ISs | 97-167 | This study, [36,<br>37] |
|  | VIM-2 (n=1) | INT_P4_C | MOB<sub>H</sub> | *Pseudomonas aeruginosa* | Class I In within Tn3-like Tn | 65 | This study |
|  | VIM-4 (n=4) | INT_P4_C | MOB<sub>H</sub> | *Pseudomonas aeruginosa*,<br>*Klebsiella pneumoniae*,<br>*Alcaligenes faecalis* | Class I In within Tn3-like Tn | 88-102 | This study |
| MPF<sub>T</sub><br>(n=16) | AFM-1 (n=1) | INT_P4_C | MOB<sub>P</sub> | *Stenotrophomonas maltophilia* | Flanked by IS91 family ISs | 77 | This study |
|  | KPC-2 (n=1) | INT_Rci_Hp1_C | MOB<sub>P</sub> | *Pseudomonas* sp. | Next to IS*Kpn6* and IS26 | 62 | This study, [6] |
|  | NDM-1 (n=3) | INT_P4_C,<br>INT_Rci_Hp1_C | MOB<sub>P</sub> | *Pseudomonas aeruginosa*,<br>*Pseudomonas asiatica* | Flanked by IS91 family ISs | 73-74 | This study, [38] |
|  | SPM-1 (n=11) | INT_Rci_Hp1_C | MOB<sub>P</sub> | *Pseudomonas aeruginosa* | Flanked by ISCR3-like elements | 44-58 | This study, [39] |
| MPF<sub>F</sub> (n=3) | IMP-6 (n=1) | DNA_BRE_C | MOB<sub>F</sub> | *Escherichia coli* | Class I In within<br>Tn3-like Tn | 118 | This study |
|  | KPC-2 (n=2) | DNA_BRE_C | MOB<sub>F</sub> | *Klebsiella pneumoniae* | Tn4401 | 75 | This study |
| Incomplete<br>MPF<sub>G</sub> (n=2) | IMP-8 (n=1) | DNA_BRE_C | MOB<sub>H</sub> | *Enterobacter cloacae* | Class I In within<br>Tn3-like Tn | 124 | This study, [40] |
|  | NDM-1 (n=1) | INT_P4_C | MOB<sub>H</sub> | *Pseudomonas aeruginosa* | Flanked by<br>ISCR3-like<br>elements | 98 | This study |
| Incomplete<br>MPF<sub>T</sub> (n=1) | KPC-2 (n=1) | INT_Rci_Hp1_C | NA | *Achromobacter* sp. | Next to ISKpn6<br>and IS26 | 43 | This study |
| Incomplete<br>MPF<sub>F</sub> (n=1) | NDM-5 (n=1) | DNA_BRE_C | NA | *Escherichia coli* | Flanked by<br>ISCR1 and IS26 | 54 | This study, [41] |

**Abbreviations:** CEG, carbapenemase-encoding gene; DNA\_BRE\_C, DNA breaking-rejoining enzymes, C-terminal catalytic domain; ICE, Integrative and Conjugative Elements; In, Integron; INT\_P4\_C, P4 integrase, C-terminal catalytic domain; INT\_Rci\_Hp1\_C, Shufflon-specific DNA recombinase Rci and Bacteriophage Hp1\_like integrase, C-terminal catalytic domain family; MPF, Mating Pair Formation; NA, no hit; ST, Sequence Type.
<sup>1</sup>Some sequences are not complete. For more details, please refer to **Table S5**.

**Figure 2.**
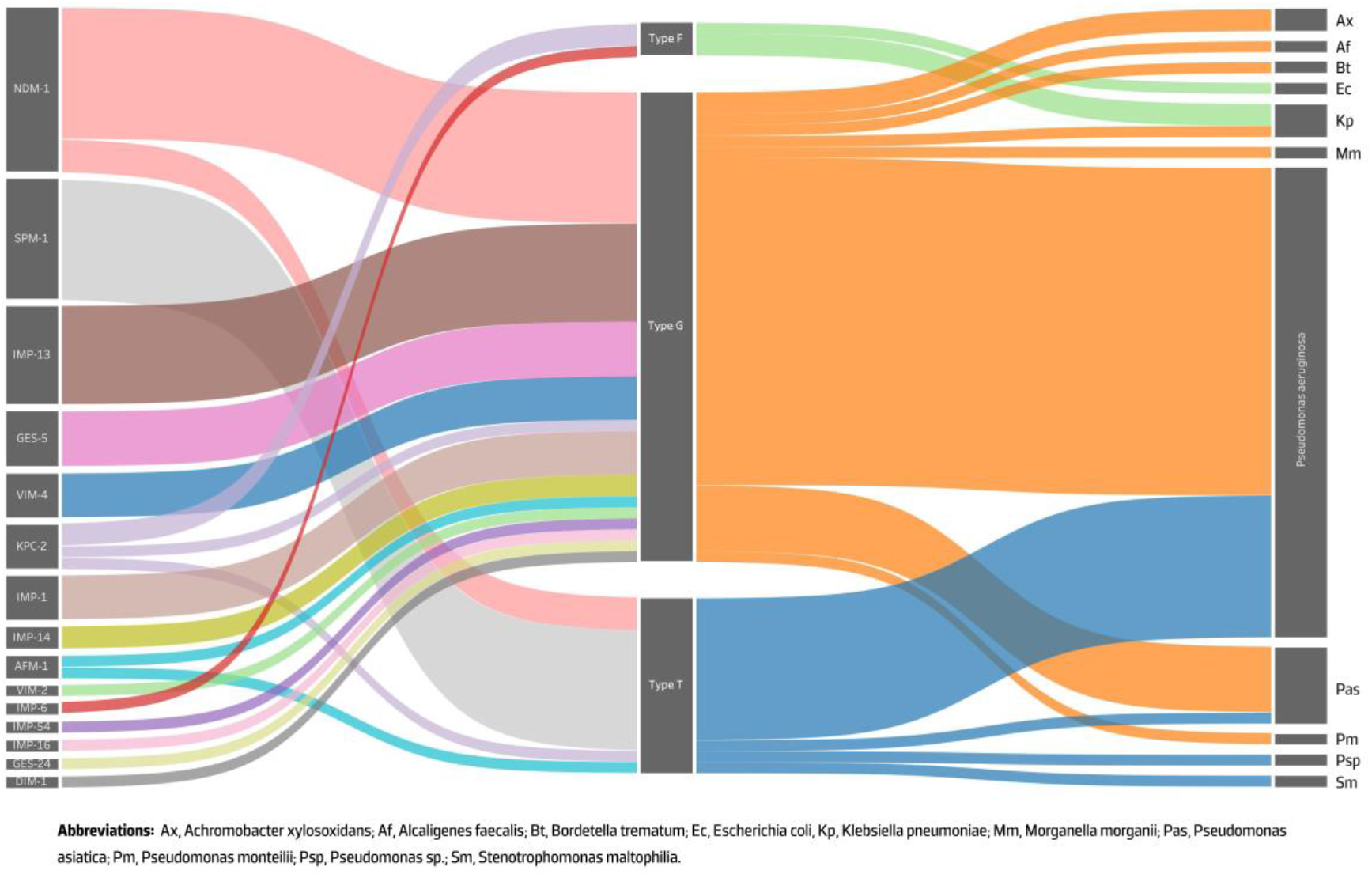
Sankey diagram showing the contribution of different MPF families for the spread of CEGs among several bacterial genomes. The left, centre and right axis represents the association between the identified carbapenemases, MPF type and the bacterial genus, respectively, while the width of each connection is proportional to the number of positive hits.

**Figure 3.**
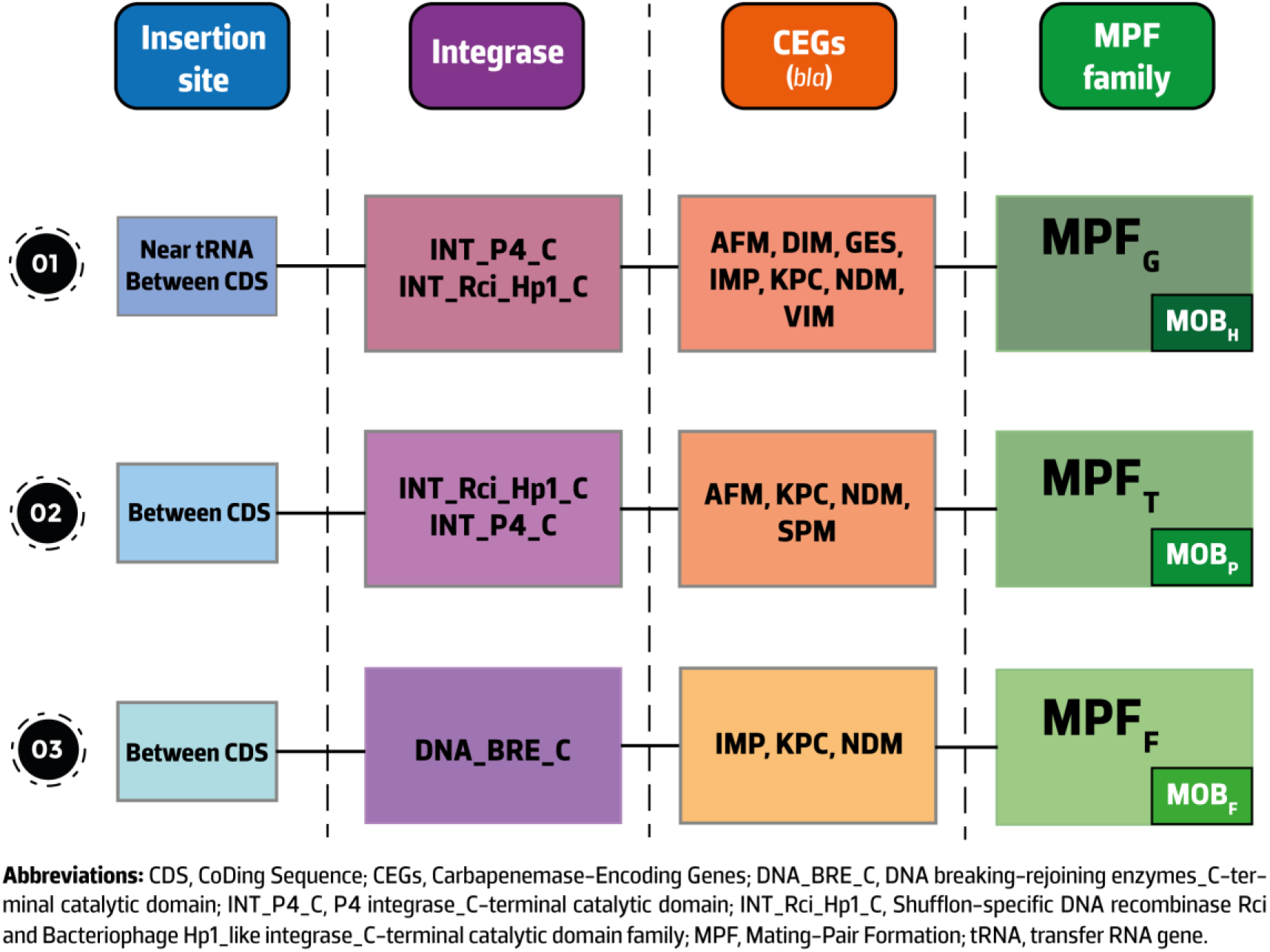
Schematic representation of the different insertion site-integrase-CEGs-MPF-relaxase profiles identified in this study. The figure is not to scale and the relative position of the different modules (integration, conjugation, accessory CEGs) is for illustrative purposes to show the various relationships observed.

Using the CONJscan module of MacSyFinder we identified the MPF family for 62 hits (incomplete MPF classes were predicted for the remaining 4 hits) and we noted that these hits belong to three families: MPF_G_ (69%, n=43/62), MPF_T_ (26%) and MPF_F_ (5%) (**Figure 2**, **Table 1**, **Table S5** and **Table S6**). In our results, the MPF_G_ class was only associated with MOB_H_, MPF_T_ with MOB_P_ and MPF_F_ with MOBF (**Figure 3**). All ICEs here identified carried a tyrosine recombinase, with the majority of them (56%, n=37/66) belonging to the P4 integrase, C-terminal catalytic domain family (INT_P4_C). The shufflon-specific DNA recombinase Rci and bacteriophage Hp1_like integrase, C-terminal catalytic domain family (INT_Rci_Hp1_C) and the DNA breaking-rejoining enzymes, C-terminal catalytic domain family (DNA_BRE_C) integrases were also identified in our collection (36% and 8%, respectively) (**Table 1** and **Table S5**). INT_P4_C and INT_Rci_Hp1_C integrases were associated with ICEs belonging to the MPF_G_ and MPF_T_ classes, while DNA_BRE_C was found on MPF_F_ ICEs (**Figure 3**). ICEs from the MPF_G_ and MPF_T_ classes were particularly promiscuous, being responsible for the spread of several CEGs of the metallo-beta-lactamase family such as *bla*_NDM-1_, *bla*_SPM-1_ and *bla*_IMP_ variants among clinically-relevant pathogens as *P. aeruginosa*, *K. pneumoniae* and *E. coli*. ICEs of the MPF_F_ class carrying blaIMP_−6_ or *bla*_KPC-2_ were restricted to *E. coli* and *K. pneumoniae*. The *bla*_SPM-1_ gene was exclusively identified in *P. aeruginosa* and in ICEs of the MPF_T_ class (**Table 1**).

We analysed the types of integrase, relaxase and MPF classes present among four model ICEs: ICE*clc*, pKLC102, SXT and Tn*4371*. The MPF_G_-INT_P4_C ICEs here identified are related to ICE*clc*, since this ICE belongs to the same class, carries a MOB_H_ relaxase and also an INT_P4_C integrase. The MPF_T_-INT_Rci_Hp1_C ICEs belong to the Tn*4371* family, which also carries the MOB_P_ relaxase. No conserved domain family could be attributed for SXT; however, the MPF_F_ ICEs here reported should be related to this model ICE that also uses a MOB_H_ relaxase.

Additionally, we also identified 386 hits encoding an integrase and a relaxase in the vicinity of CEGs (**Table S7**). For these hits, however, we could not predict if the CEG is located on a plasmid or an ICE, since the contig is fragmented and tracing the boundaries of the element is not possible, or the sequences have assembly gaps that make this prediction challenging. Also, some plasmids may encode a tyrosine or serine recombinase, and some ICEs may encode replicases and partition systems that are typical of plasmids [34], which can hinder the accurate prediction of genetic platforms when the sequence has poor quality or is highly fragmented due to short-read sequencing approaches.

### A variable repertoire of CEG-bearing integrons and transposons target ICEs

We identified 17 CEG variants among the 66 putative ICEs, dominated by *bla*_NDM-1_ and *bla*_SPM-1_ (**Table 1**). Insertion sequences (ISs, e.g. ISCR3-like elements) were frequently linked to the acquisition of *bla*_SPM-1_ and *bla*_NDM-1_ [38, 39], while *bla*_IMP_, *bla*_VIM_, and *bla*_GES_ were found on class I integrons frequently integrated into Tn*3*-family transposons [6]. The *bla*_KPC-2_ gene was typically found within Tn*4401*-like transposons, which are capable of conferring a high frequency of transposition [42]. The recently identified AFM-1 metallo-beta-lactamase (Genbank accession number MK143105.1) was here identified in two ICEs inserted in *Bordetella trematum* and *Stenotrophomonas maltophilia* genomes (**Table 1**). We also found *bla*_NDM-1_ genes in ICEs integrated into the genomes of a recently proposed *Pseudomonas asiatica* species, which is spreading in hospital settings in Myanmar [43]. Besides CEGs, the ICEs identified in this study also harbour genes conferring resistance to other antibiotics, such as aminoglycosides, fluoroquinolones, macrolides, and tetracyclines (**Table S8**), widening the spectrum of transmissible AR genes selectable by carbapenems due to linkage.

### Acquisition of additional traits by ICEs include competitive weapons as bacteriocins and siderophores

Besides genes conferring antibiotic resistance, the CEG-bearing ICEs here identified harbour other cargo genes that may confer a selective advantage to the ICE host. We found DUF692 domains typical of bacteriocin production genes among the 6 MPF_G_ ICEs from *P. asiatica* strains and the *P. aeruginosa* N15-01092, 1334/14 and ST773 strains (**Table S5**). All these ICEs carry a *bla*_NDM-1_ gene and a similar copy of the bacteriocin-encoding gene. Curiously, the bacteriocin production gene was not identified in the MPF_T_ ICE from *P. asiatica* strain MY569. Additionally, we found the siderophore aerobactin operon within an ICE in *E. coli* strain E302 (**Table S5**). This operon is usually found in enterobacterial plasmids, and was also identified in a pathogenicity island in uropathogenic *E. coli* strain CFT073 [44]. We identified no CRISPR-Cas systems among the ICEs here identified. Nearly half of them (47.0%, n=31/66) carried complete or incomplete restriction-modification systems belonging to type II, III and IV (**Table S9**).

The majority of the proteins encoded within the 66 ICEs refer to replication, recombination, transcription and intracellular trafficking functions (**Figure 4**). Several proteins, however, encoded for unknown functions (34.0%, n=1,615/4,754, **Table S10**), highlighting the lack of knowledge on the ICE proteome.

**Figure 4.**
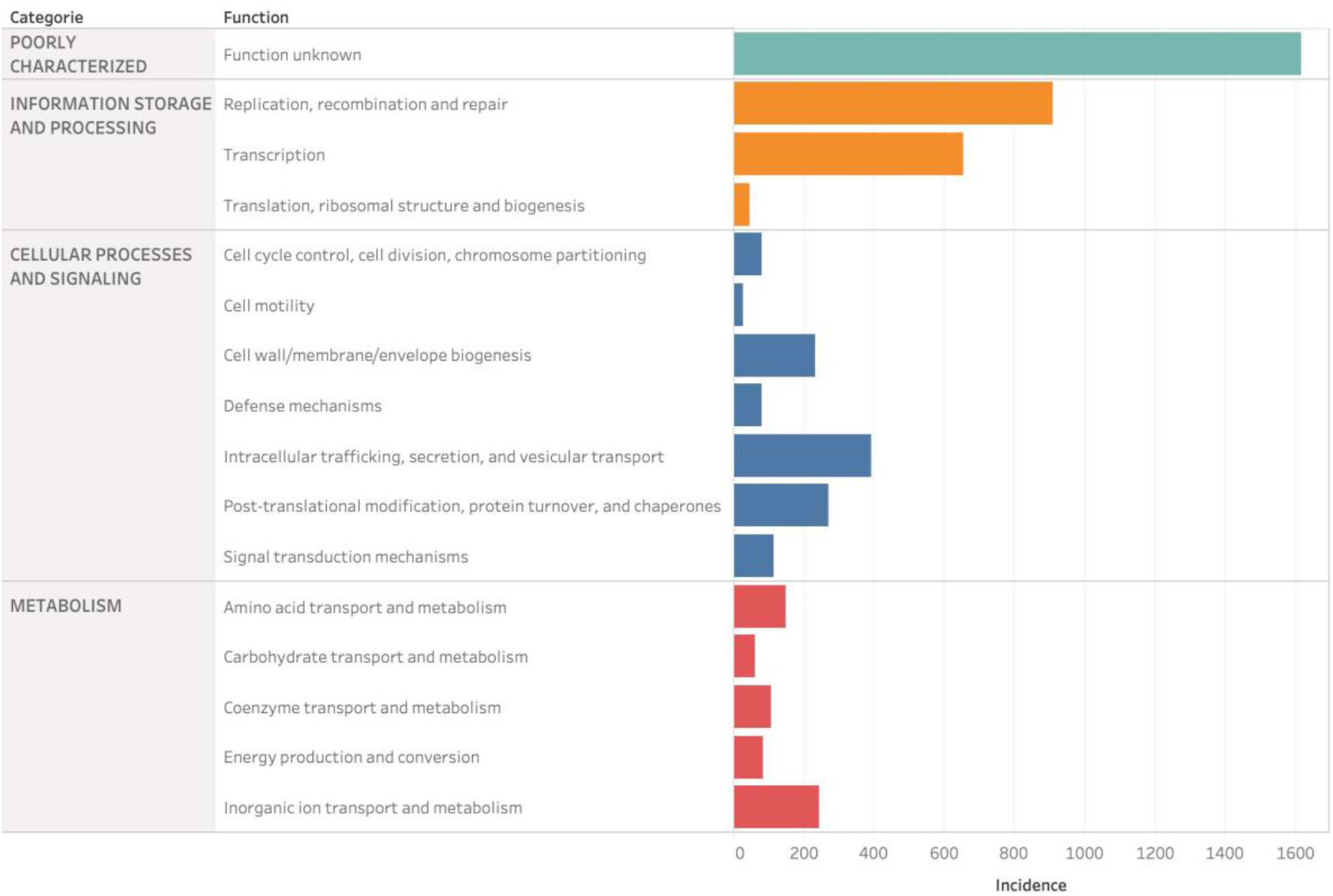
Grouped bar chart representing the incidence of each eggNOG function broken down by category (represented by different colours) in the CEG-bearing ICE sequences.

## Discussion

We have set out to comprehensively identify the CEGs among all bacterial genomes deposited in NCBI and the CEG-bearing ICE sequences. Our study considerably expands the knowledge of the repertoire of CEGs-bearing ICEs. We uncovered 66 putative ICEs that may be involved in HGT of CEGs amongst bacterial genomes. To expand our predictions, we also used the CONJscan module of Macsyfinder to trace the MPF families likely involved in HGT. Our analysis on the co-occurrence of relaxases with MPF families (**Table 1** and **Table S5**) is in agreement with the combinations observed by Guglielmini and colleagues [11]; all ICEs belonging to the MPF_G_ class carried a MOB_H_ relaxase, and the MOB_P_ and MOB_F_ relaxases were linked to MPF_T_ and MPF_F_, respectively. All MPF_G_ class ICEs here described present a MPF class-relaxase-integrase profile that resembles that from ICE*clc*. Even though pKLC102 is also a representative of the MPF_G_ class and carries a MOB_H_ relaxase, it uses a DNA_BRE_C integrase instead of a INT_P4_C. The absence of CEG-carrying ICEs from the pKLC102 family was already reported [6].

The scenario observed for the acquisition of the most important CEGs by ICEs (ISs for *bla*_NDM-1_ and *bla*_SPM-1_, class I integrons for *bla*_IMP_ and *bla*_VIM_ and Tn*4401* for *bla*_KPC-2_) resembles that of plasmids [3] and provide additional support to the notion that the line separating these elements is blurred [13, 14]. We now show that besides plasmids, this promiscuous repertoire of integrons and transposons frequently targets ICEs of different MPF families. Even though CEGs might rapidly spread worldwide, local selection is likely required for them to reach fixation, as can be seen for the clonal expansion of *P. aeruginosa* ST277 harbouring *bla*_SPM-1_ [39]. Surprisingly, we noted that this gene was not detected beyond the same clonal lineage. Indeed, all hits were identified in *P. aeruginosa* ST277 strains from Brazil and within a MPF_T_ ICE family, indicating that these ICEs are transferring vertically and/or horizontally within this ST. Understanding the limitations on HGT of this family of ICE may be translatable to other, more transferable, ICEs and could underpin a control strategy to prevent the spread of these elements in the future.

Although *bla*_KPC_ and *bla*_OXA_ were the most frequently identified CEGs within the analysed genomes (**Table S2**), we only found 5 ICEs carrying *bla*_KPC-2_ and 2 ICEs with *bla*_OXA_ genes (**Table S8**). The *bla*_KPC_ genes are mostly located in Tn*4401* transposons that target *K. pneumoniae* plasmids, while the *bla*_OXA_ genes are frequently associated with ISs that tend to target *Acinetobacter* spp. chromosomes and plasmids [3]. These genes may take advantage of the copy number and the higher genetic plasticity of plasmids [34]. This plasticity may increase the rate of appearance of novel mutations and the high copy number may amplify the effect due to the increased gene dosage [45].

In addition to AR, ICEs can be involved in other adaptive traits such as carbon-source utilization, symbiosis, restriction-modification, siderophore and bacteriocin synthesis [14]. Since bacteria commonly inhabit highly competitive environments, the production of specific secondary metabolites (such as bacteriocins and siderophores) may confer a selective advantage to the host [46]. We speculate that the presence of these metabolites within the ICEs here characterized may promote their stability by preferentially selecting for cells harbouring the ICE. CRISPR-Cas systems are rarely found on MGEs, and the type I-C systems carried within pKLC102-related ICEs are one of these few examples [9]. Since none of the ICEs here described belong to the pKLC102 family, the absence of CRISPR-Cas systems within our dataset was expected.

One caveat of our studies is that these results do not yet expose the complete set of CEG-bearing ICEs present in all bacteria. There is an inherent bias in the number of times we detect a particular CEG in certain bacterial genomes as some are over represented in the database compared to others. It is possible that certain observations will flatten out as more genomes are analysed. Less than 10% of the bacterial genomes currently present on NCBI are complete. This is a major drawback since putative ICEs present in draft genomes tend to be fragmented due to the presence of repetitive regions that are not resolved using short-read sequencing. Also, the relaxase and the T4SS encoded by ICEs resemble those of plasmids [11, 13, 47]. Plus, it is possible that ICEs and plasmids have swapped conjugation modules throughout their evolutionary history [11]. We believe that a more thorough exploration of this issue, especially regarding the precise delimitation of ICEs, will be an important further step toward an improved understanding of the contribution of these elements to bacterial adaptation and evolution of AR.

While we have chosen to focus on CEG-bearing genomes, our computational approach can be applied to trace other relevant AR genes and other cargo genes that may confer a selective advantage to the ICE host. Leveraging knowledge linking the accurate prediction of ICE sequences to the carriage of AR genes, will not only improve our understanding of HGT, but may also uncover potential approaches to tackle the spread of AR.

## Abbreviations

AR: antibiotic resistance
CEG: carbapenemase-encoding gene
HGT: horizontal gene transfer
ICE: integrative and conjugative element
MGE: mobile genetic element
MPF: mating pair formation
NCBI: National Center for Biotechnology Information
T4SS: Type-IV Secretion System
WHO: World Health Organization

## Data bibliography

The DNA sequences used in this study are available in NCBI, under the accession numbers provided in supplementary tables S1-S9.

## Funding information

This work was supported by the Applied Molecular Biosciences Unit - UCIBIO which is financed by national funds from FCT (UIDB/04378/2020).

## Author contributions

J. B. conceptualized the project. J. B. ran the analyses. J.B. and J. M. analysed the data. J.B. wrote the manuscript. J. M., A. P. R. and L. P. edited and revised the manuscript. All authors read, commented on and approved the final manuscript.

## Conflicts of interest

The authors declare that there are no conflicts of interest.

## Notes

https://figshare.com/projects/_Comprehensive_genome_data_analysis_establishes_a_triple_whammy_of_carbapenemases_ICEs_and_multiple_clinically-relevant_bacteria/78369

### Competing Interest Statement

The authors have declared no competing interest.

